# Inhibiting fibroblast aggregation in skin wounds unlocks developmental pathway to regeneration

**DOI:** 10.1101/608075

**Authors:** Ashley W. Seifert, Adam B. Cook, Douglas Shaw

**Author notes:** To whom correspondence should be addressed: Ashley W. Seifert.

## Abstract

Salamanders are capable of full-thickness skin regeneration where removal of epidermis, dermis and hypodermis results in scar-free repair. What remains unclear is whether regeneration of these tissues recapitulates the cellular events of skin development or occurs through a process unique to regenerative healing. Unfortunately, information on the post-embryonic development of salamander skin is severely lacking, having focused on compartments or cell types, but never on the skin as a complete organ. By examining coordinated development of the epidermis and dermis in axolotls we establish six distinct stages of skin development (I-VI): I-V for normally paedomorphic adults and a sixth stage following metamorphosis. Raising animals either in isolation (zero density pressure) or in groups (density pressure) we find that skin development progresses as a function of animal size and that density directly effects developmental rate. Using keratins, p63, and proliferative markers, we show that although the epidermis lacks visible stratification at early stages of skin development, when the dermis transforms into the stratum spongiosum and stratum compactum keratinocytes differentiate into at least three distinct phenotypes that reveal a cryptic stratification program uncoupled from metamorphosis. Lastly, comparing skin regeneration to skin development, we find that dermal regeneration occurs through a unique process, relying heavily on remodeling of the wound extracellular matrix, rather than proceeding through direct development of a dermal lamella produced by the epidermis. By preventing fibroblast influx into the wound bed using beryllium nitrate, we show that in the absence of fibroblast generated ECM production skin regeneration occurs through an alternate route that recapitulates development.

## INTRODUCTION

Urodeles have long served as model organisms for developmental and regenerative biology (Callery, 2006). Major contributions to developmental biology concerning the inductive properties of the organizer, regulative development of the limb field, and the regulation of organ size, were made using salamanders and newts. Similarly, these animals have played an outsized role in the field of regenerative biology because of their extensive powers of organ regeneration: adult salamanders and newts can regenerate almost any organ or tissue in the body including the spinal cord, brain, limb and skin (Carlson, 2007). It seems natural then, that similarities between embryonic development and regeneration would contribute to studies investigating the mechanistic basis for how tissues are rebuilt after damage. An oft-invoked hypothesis is that vertebrate regenerators re-deploy developmental programs to orchestrate regeneration (i.e., organ regeneration recapitulates organ development). Despite the fact that regeneration takes place in an adult environment and involves processes that do not occur during embryonic development (e.g., wound healing, inflammatory and immune cell influx, etc.), this hypothesis has important implications for understanding how resident cells orchestrate tissue regeneration as the injury microenvironment transitions towards rebuilding missing tissue.

Beyond serving as a simple barrier against the external environment, vertebrate skin displays a wide range of evolutionary adaptations to specific environments. As a protective barrier it is frequently injured and, depending on the species, either regenerates the dermis, epidermis and associated appendages or forms a dense collagen scar to repair the injury (Seifert and Maden, 2014). The basic anatomy of vertebrate skin is generally conserved: there is an outer layer (epidermis) which is variably stratified and gives rise to epithelial appendages (e.g., hair, feathers, glands, etc.) and an inner layer (the dermis) comprising a network of connective tissue embedded with nerves, blood vessels and the epithelial derived appendages. Despite a vast trove of literature describing the form and function of adult skin across a panoply of species, our knowledge of skin development is noticeably uneven. Skin development has been most extensively studied in mammals, where the cellular and molecular understanding of epidermal stratification (rev in. Koster and Roop, 2007) and hair follicle formation (rev. in Schmidt Ullrich and Paus, 2005) has far outpaced our knowledge of dermal development (Driskell et al., 2013; Van Exan and Hardy, 1984). Renewed interest into the scar-free healing ability of salamanders (Cook and Seifert, 2016; Erickson and Echeverri, 2018; Huang et al., 2017; Levesque et al., 2010; Seifert et al., 2012b; Weiss and Ferris, 1956) and frogs (Bertolotti et al., 2013; Otsuka-Yamaguchi et al., 2017; Seifert and Maden, 2014; Yokoyama et al., 2011b) has exposed a relatively sparse literature on skin development in urodeles (Gerling et al., 2012; Hay and Revel, 1963; Pederzoli et al., 2002; Salpeter and Singer, 1959; Weiss and Ferris, 1954). Instead, most of our knowledge of amphibian skin development comes from work in anurans (rev. in Fox, 1986a; Fox, 1986b). Studies have detailed the progressive transformation of the epidermis and dermis leading up to and through metamorphosis (Fox and Whitear, 1990; Kawai et al., 1994; Kemp, 1959; Robinson and Heintzelman, 1987; Takagi, 1956; Tamakoshi et al., 1998) and how different keratinocyte populations contribute to epidermal stratification (Izutsu et al., 1993; Kinoshita and Sasaki, 1994; Nishikawa et al., 1992; Suzuki et al., 2002; Watanabe et al., 2002). Anuran skin development is directly controlled by thyroid hormone which acts to regulate maturation of the epidermis and dermis separately (Brown, 1997; Schreiber and Brown, 2003). Neotenic salamanders like the axolotl do not normally undergo metamorphosis and thus the cellular and molecular details of skin development in these animals remains poorly understood.

Despite a few known exceptions (e.g., African spiny mice and rabbits) (Breedis, 1954; Jiang et al., 2019; Seifert et al., 2012a), most adult mammals repair full-thickness skin wounds with scar tissue (Clark, 1996). In contrast, fetal mammals can regenerate skin wounds until late gestation (Adzick and Longaker, 1992) and studies in other vertebrate models of regeneration have demonstrated full-thickness skin regeneration in adult fish (Richardson et al., 2013), salamanders (Cook and Seifert, 2016; Levesque et al., 2010; Seifert et al., 2012b), and frogs (Bertolotti et al., 2013; Otsuka-Yamaguchi et al., 2017; Seifert and Maden, 2014; Yokoyama et al., 2011a). When wounded, the axolotl (*Ambystoma mexicanum*) undergoes a regenerative process which allows it to completely repair full-thickness skin through remodeling of fibrotic connective tissue (Cook and Seifert, 2016; Levesque et al., 2010; Seifert et al., 2012b). Previous work in aquatic and terrestrial forms noted rapid re-epithelialization, inflammatory cell influx, production of an extracellular matrix (ECM) high in tenascin-C, increased matrix metalloproteinase expression, remodeling of the ECM and replacement of missing skin layers (Seifert et al., 2012b). Interestingly, although axolotls heal scar-free, an extensive cross-species comparison with other regenerators, and with fibrotic repair in mammals, supports a model where fibrosis generates excessive ECM that is either remodeled into scar tissue or remodeled into a facsimile of the original tissue (Seifert and Maden, 2014). This model suggests that although similarities between development and regeneration may exist, the two processes are perhaps more distinct than often appreciated. In fact, early descriptions of amphibian epidermis and basement lamella development (Hay and Revel, 1963; Kemp, 1961; Utoh et al., 2000; Weiss and Ferris, 1954) suggest that remodeling of a dense fibrotic wound matrix into a bi-layered dermis may be a process distinctly different from development.

Here we investigate post-embryonic skin development and regeneration in axolotls. By examining distinct cellular features of the epidermis and dermis we establish six distinct stages of skin development (I-VI): I-V for normally paedomorphic adults and a sixth stage following metamorphosis. Raising animals either in isolation (zero density pressure) or in groups (density pressure) we find that skin development progresses as a function of animal size and that density directly effects developmental rate. Using keratins, p63 and proliferative markers we show that the epidermis lacks visible stratification at early stages of skin development. However, when the dermis begins to mature into the stratum spongiosum and stratum compactum, keratinocytes differentiate into at least three distinct phenotypes that reveal a cryptic stratification. Lastly, comparing skin regeneration to skin development, we find that despite some similarities, dermal regeneration occurs uniquely, relying heavily on remodeling of the wound extracellular matrix from the margins, rather than proceeding through developmental events. By preventing fibroblast influx into the wound bed using beryllium nitrate, we show that delayed skin regeneration observed in these animals proceeds through an alternate route that recapitulates development.

## RESULTS

### Axolotl skin progresses through six developmental phases

In order to understand how ectoderm and mesoderm undergo morphogenesis to form the skin in salamanders, we analyzed developing flank skin in the axolotl *Ambystoma mexicanum* (Fig. 1A-F). First, we sampled skin from post-hatching animals across a broad range of ages and sizes to identify consistent cellular features that we could use to establish a regular and repeated series of developmental skin stages. Using this approach, we defined five distinct phases through which the skin transitions during post-embryonic development into its stable adult phenotype and an additional transition following induced metamorphosis (Table 1 and Fig. 1A-F).

**Figure 1.**
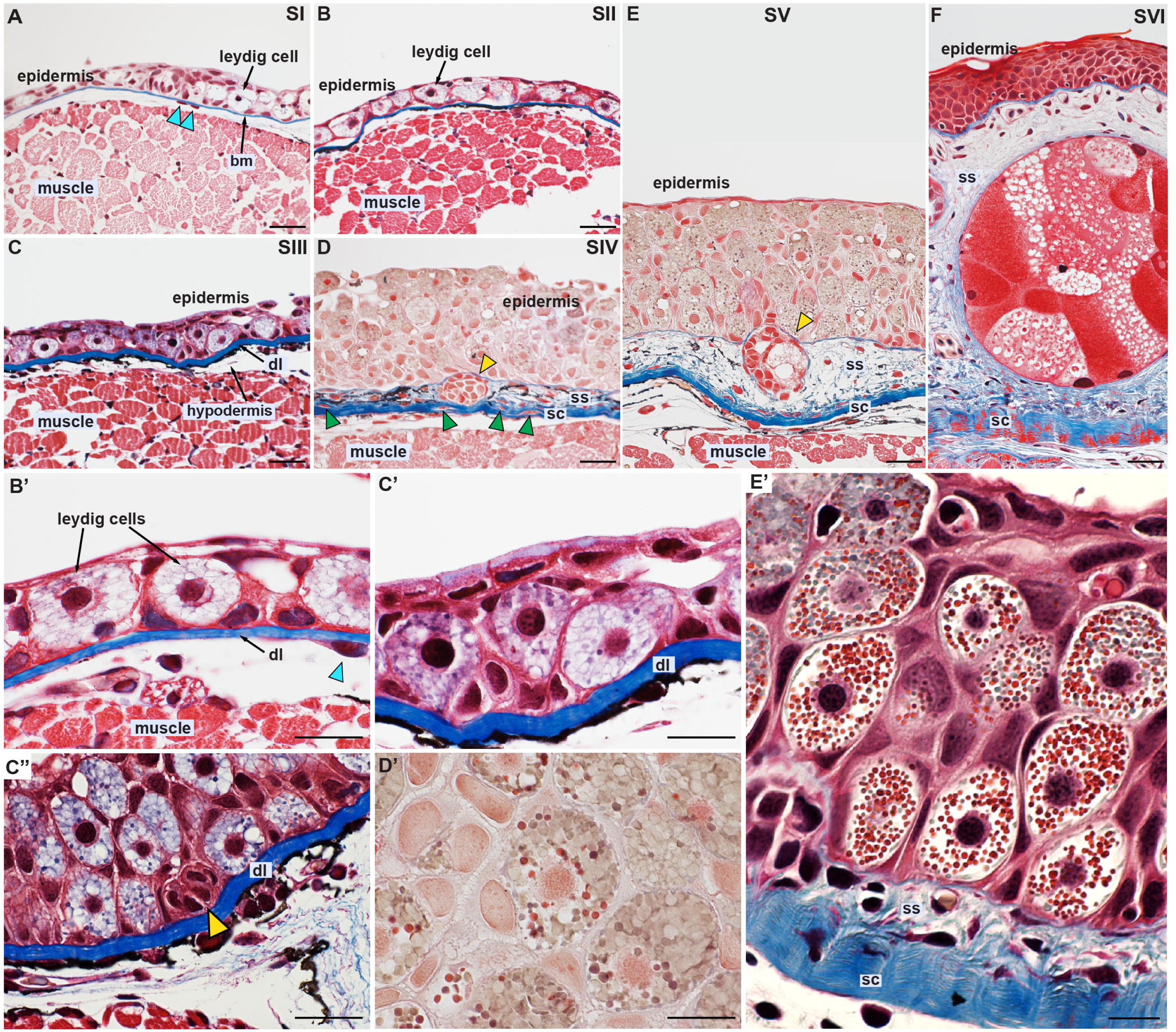
Axolotl skin development can be categorized into six distinct stages. (A-F) Histological skin preparations stained with Masson’s Trichrome depicting developmental stages I-VI (see Table 1 for detailed description of each stage). Each panel is representative of the indicated stage. All samples were obtained from the dorsal flank region, posterior to the front limb. Extracellular matrix/dermal components appear blue, cellular peptides/nucleus stained purple, muscle stained red. (A) Leydig cells appear in the epidermis at stage I along with a prominent basement membrane. Mesenchymal cells are found beneath the basement membrane (blue arrows). (B’-E’) Leydig cell differentiation is associated with increasing granulation around the nucleus. (B-B’) Leydig cells occupy the majority of the epidermis in stage II skin and show vacuoles around the nucleus without granules. The dermal lamella is first visible as a slight thickening in association with the basement membrane. (C’-C”) At stage III, keratinocyte numbers increase in the epidermis and granules appear within Leydig cells as the epidermis thickens into a multi-layered structure. The acellular dermal lamella thickens in association with the basement membrane and forming glands (yellow arrow) are visible in the epidermis (C”). (D-D’) Leydig cell granulation increases in stage IV skin as the epidermis transforms from a simple to transitional epithelium. The dermal lamella delaminates from the basement membrane as fibroblasts (green arrows) and melanocytes invade the dermal lamella (D). Glands (yellow arrow) descend into the dermis as it begins to stratify. (E’) Leydig cells are fully differentiated in stage V skin which represents the adult phenotype in paedomorphic adults. (F) Stage VI occurs after induced metamorphosis. Leydig cells completely disappear from the epidermis which transforms into a fully stratified epithelium. Granular glands enlarge to occupy the entire stratum spongiosum (ss). BM = basement membrane, DL = dermal lamella, SS = stratum spongiosum, SC = stratum compactum. Scale bars = 50μm (A-F) and 20μm (B’-E’).

**Table 1.** Developmental stages of skin development in the axolotl, *Ambystoma mexicanum*. Stages were determined post-hoc after histological characterization of unique structural and cellular changes in the epidermis and dermis. These included: proportion of Leydig cells and keratinocytes in epidermis, degree of Leydig cell differentiation (amount of granulation), epidermal and dermal stratification, epidermal/dermal thickness and gland development. See Figure 1 for cellular representation of stages.

| Skin Development Stage | Diagnostic Features |
| --- | --- |
| Stage I | Epidermis comprised primarily of epithelial cells and some vacuolated Leydig cells. Thin basement membrane present beneath epidermis. No more than two cell layers thick. No dermis. |
| Stage II | Epidermis comprised primarily of Leydig cells. Granules begin to appear in Leydig cells. Thin basement membrane present beneath epidermis. First appearance of dermal lamella is association with basement membrane. No more than two cell layers thick. |
| Stage III | Epidermis several layers with mixture of keratinocytes and Leydig cells. Thickening of dermal lamella. Keratinocytes begin aggregating to form glands. |
| Stage IV | Epidermis resembles transitional epithelium. Dermis begins stratifying into stratum spongiosum and stratum compactum. Cells invade dermis and immature glands begin descending into stratum spongiosum. Leydig cells appear fully differentiated. |
| Stage V | Epidermis remains transitional epithelium. Mature glands in stratum spongiosum. Mature, bi-layered dermis with cellular stratum spongiosum and thickened acellular stratum compactum. |
| Stage VI | Following metamorphosis leydig cells disappear from the epidermis. The epidermis transforms from a transitional epithelium to a stratified epithelium. Granular and mucous glands in the dermis enlarge to fill most of the stratum spongiosum. |

Amphibian pre-hatch embryos possess a single layer of ectodermally-derived epithelial cells that surround the embryo atop a thin membrane separating the epidermis from the underlying muscle. After hatching, the epidermis contained multiple layers of epithelial cells that were interspersed with Leydig cells and we identified this as the first post-embryonic stage of skin development (stage I) (Fig. 1A and Table 1). In stage I skin, Leydig cells exhibited the early stages of differentiation with a single nucleus surrounded by large vacuoles (Fig. 1A) (Gerling et al., 2012). Ampullae associated with the developing lateral line were also visible in the epidermis and cells within the ampullae were positive for keratin expression (Fig. 1A, asterisk and Supplementary Fig. 1A-D). A dermal layer was absent at stage I and the interstitial space beneath the basement membrane was sparsely populated with mesenchymal cells (Fig. 1A). As development proceeded, the skin transitioned through a phase (stage II) characterized by subtle and distinct changes in the epidermis (Fig. 1B-B’). Specifically, Leydig cells now occupied the majority of the epidermis and small granules began to appear within the vacuolated spaces close to their nuclei (Fig. 1B-B’). Additionally, a thickened band of collagen, the dermal lamella, first appeared beneath the epidermis (Fig. 1B-B’). The next phase of skin development (stage III) was marked by a prominent thickening of the dermal lamella contiguous with the basement membrane and an increase in the number of epithelial cells within the epidermis (Fig. 1C-C’). Within the epidermis at this stage groups of keratin-positive cells were observed aggregating to form the anlagen of glands (Fig. 1C” and Suppl. Fig. 1E). We also observed an increase in pigment and mesenchymal cells throughout the hypodermis beneath the dermal lamella (Fig. 1C-C”). No mesenchymal cells were found within or above the dermal lamella. Stage IV represented a major transformation of the epidermis and dermis (Table 1 and Fig. 1D-D’). During this phase, the epidermis matured into a transitional epithelium (similar to the mammalian urinary, ureter and bladder epithelium) and more than doubled in thickness as the number of epithelial and Leydig cells increased (Fig. 1D). Leydig cells in Stage 4 skin were now differentiated with varying amounts of granules tightly surrounding the nucleus (Fig. 1D-D’). In parallel with epidermal maturation, fibroblasts and pigment cells invaded into and above the dermal lamella as it separated from the epidermis (Fig. 1D and Supplementary Fig. 1C). We also observed glands descending through the basement membrane as the stratum spongiosum began forming and the dermal lamella became the stratum compactum (Fig. 1D). During the final phase of skin development in paedomorphic axolotls (stage V), the dermis completed its stratification and descendant glands in the stratum spongiosum differentiated into granular or mucous glands, the two primary glands present in salamander skin (Fig. 1E). Leydig cells continued to occupy a majority of the epidermis which remained a transitional epithelium and the stratum corneum was visible as a thin layer of flattened keratinocytes covering the outermost layer of the epidermis (Fig. 1E).

Captive, wildtype axolotls do not normally undergo metamorphosis and thus, stage V is the normal adult phenotype in paedomorphic animals. However, axolotls do retain the ability to undergo metamorphosis and we induced this transformation using thyroxine (Coots and Seifert, 2015) which caused the skin to transition through a sixth phase (stage VI) (Table 1 and Fig. 1F). During this transition, the epidermis transformed into a stratified epithelium and Leydig cells completely disappeared (Fig. 1F). The stratum compactum persisted as a thickened collagen band and remained separated from the flank muscle by the hypodermis. In response to thyroxine, granular glands increased in size to occupy most of the stratum spongiosum (Fig. 1F). Together, these data provide a classification scheme defining six transitional phases of skin development in axolotls, with the post-metamorphic phase representing an adult phenotype similar to other terrestrial urodeles and anurans.

### Progression of skin development is driven by organismal size

Previous work in anurans found considerable variability during the early phases of skin development among similar aged animals (Kawai et al., 1994) and we noticed that despite being raised together, some same-aged (post hatch) animals were at different stages of skin development. Although natural populations of salamanders experience density-dependent effects on growth rate, post larval development and metamorphosis (Semlitsch, 1987), the degree to which density directly effects skin development is unknown. To directly address this question, we conducted a series of experiments to establish how skin development progresses as a function of size (mass and length) and age (post-hatch) under zero density pressure (animals raised in isolation) and density pressure (animals raised in groups). First, we individually raised sibling axolotls (n=31) immediately after hatching while regulating temperature, light, and food intake. At regular 1-week intervals we measured length (snout-tail length, cm) and mass (g), and then randomly selected individual animals in order to histologically prepare dorsal flank skin samples. When analyzing sections to determine the phase of skin development we remained blind to animal character information: mass, length, and age. Running regression analyses using stage as the response variable and mass, length, and age as predictor variables, we found that all three parameters were highly correlated with stage of skin development (adjusted r^2^ mass = 0.959, *F*= 257.708, *P*< 0.0001; adjusted r^2^ length = 0.954, *F*= 228.891, *P*< 0.0001; adjusted r^2^ age = 0.855, *F*= 65.690, *P*< 0.0001) (Fig. 2A). In principle, using regressions for any of these parameters should identify skin development stage for a particular animal raised under different conditions.

**Figure 2.**
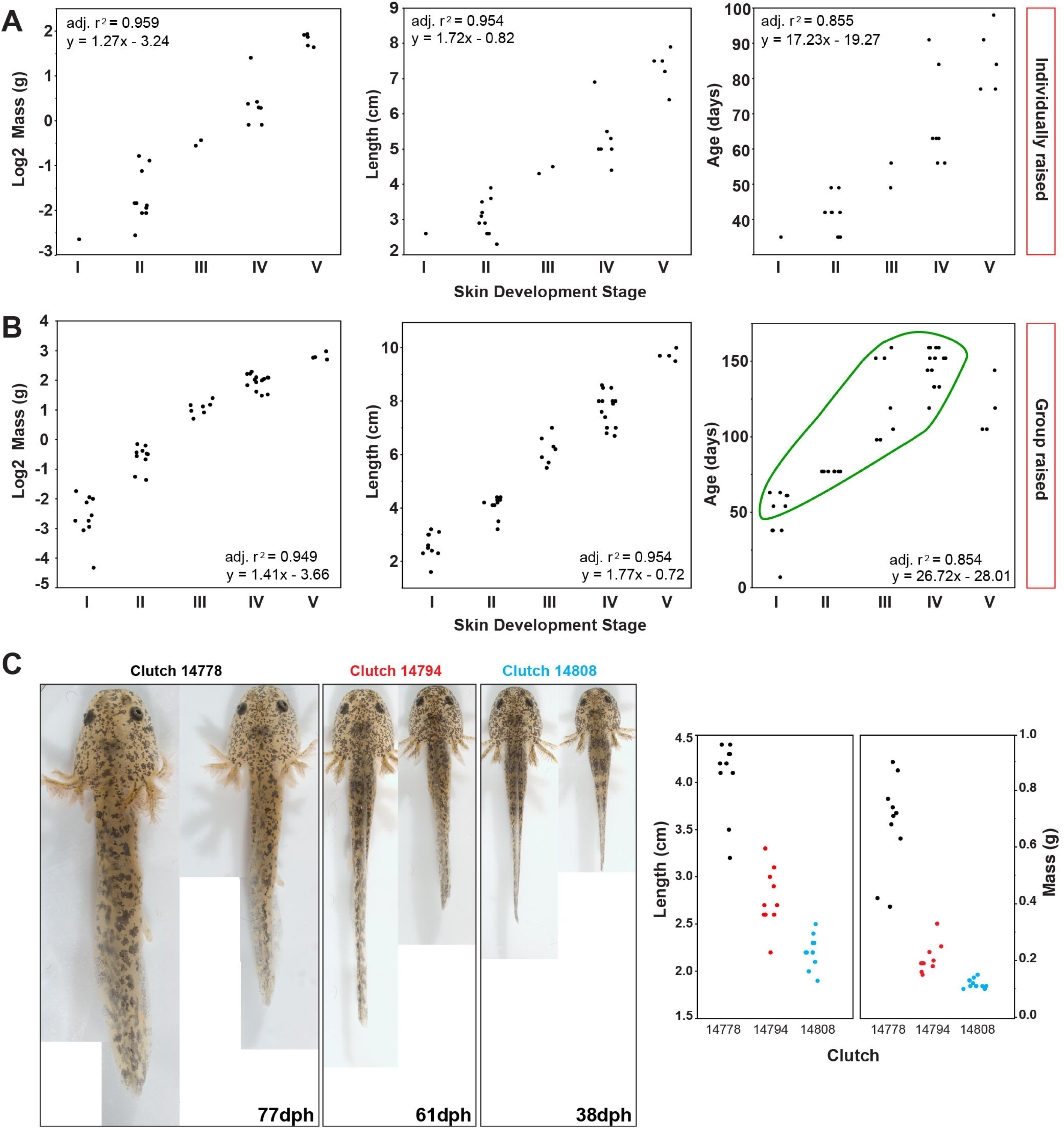
Skin development is strongly correlated with animal size and density effects developmental rate. (A-B) Animal mass, total length, and age plotted as a function of skin stage (I-V) for individually (A) raised (n=31) and group raised (n=45) post-hatching axolotls (B). Mass and length show the strongest correlation with developmental stage for zero-density and density conditions. Density reduced growth rate (B) and when skin development stage was estimated based on regressions calculated for mass, length and age from individually raised animals, 82% of the predictions overestimated their histologically-determined stages (green circle in age plot). (C) Group-raised animals from the same clutch paired to show size disparity within the same age group. Size variance within clutches increased as a function of age from the same clutch (black = 77dph, red = 61dph, blue = 38dph). Data used to generate regressions available as Supplementary Data Table 1.

In order to test the predictive power of these regressions and whether density effected developmental timing, we characterized skin developmental stage in group-raised animals (Fig. 2B) (see Methods for details). We harvested animals (n=45) from seventeen clutches across a range of ages, measured mass and length, and then used the regressions calculated from individually-raised animals to predict skin development stage. Next, we histologically prepared flank skin samples from selected animals and compared the actual stage of skin development to our predicted stages. When analyzing sections to determine the phase of skin development we again remained blind to animal character information (mass, length, and age). We found that mass and length accurately predicted skin stage 91% (41/45) and 89% (40/45) of the time respectively, whereas age was only 18% (8/45) accurate in predicting developmental skin stage (Fig. 2B). Observing multiple animals from individual clutches, it was clear that density produced considerable size variation among same-aged individuals (Fig. 2C). In some cases, clutches that differed by as much as three weeks post hatch contained individuals of similar sizes (Fig. 2C). Thus, most of the error in accurately identifying developmental skin stage for a particular animal occurred because of density driven reduction in growth rate compared to animals raised in isolation (Fig. 2B, green circle). Taken together, our data demonstrates that skin development progresses as a function of animal size and that increased density affects skin development rate by slowing growth rates.

### Keratinocyte proliferation occurs evenly throughout the epidermis until metamorphosis when it becomes restricted to basal cells

Having established the normal progression of skin development in salamanders, we next sought to molecularly characterize cellular differentiation of the epidermis. First, we examined the distribution of proliferating cells in the epidermis beginning at stage 1. Using an antibody to detect PCNA we found that proliferating (PCNA+) keratinocytes were distributed evenly across the entire epidermis with no apparent regionalization (Fig. 3A-A’). We also assessed localization of p63 which is normally restricted to basal keratinocytes in other terrestrial vertebrates (Truong et al., 2006). Echoing the distribution of PCNA, p63+ keratinocytes were found all throughout the epidermis (Fig. 3A”). While the same even distribution of PCNA+ and p63+ keratinocytes was observed through stage 5, the stratum corneum rarely contained p63+ cells after stage 2 (Fig. 3A-E”). After metamorphosis, however, we found that PCNA+ and p63+ keratinocytes were almost exclusively restricted to the basal epithelial layer (Fig. 3F-F). Suprabasal keratinocytes immediately above the basal layer also were PCNA+ and p63+, but expression was reduced compared to basal keratinocytes (Fig. 3F-F”). Interestingly, we also observed that a pan cytokeratin antibody that recognizes a wide spectrum of type 1 and type 2 keratins was strongly localized to basal keratinocytes in metamorphic skin (Fig. 3F-F’). To more explicitly determine if there was regional variation among keratinocyte populations within the epidermis, we investigated the localization of keratin 5 & 10; proteins which are normally expressed in the basal and spinous layers respectively in mammals (Byrne et al., 1994). Surprisingly, we found that keratin 5 was expressed in all keratinocytes through stage V, similar to the pattern we observed for pan cytokeratin (Fig. 4A-E). Although in metamorphic skin keratin 5 localization was biased to the basal keratinocytes, it was also expressed in the spinous and granular layers (Fig. 4F). Prior to expansion of the epidermis at stage III, keratin 10 was expressed by all keratinocytes and leydig cells at stages I-II similar to keratin 5 (Fig. 4A-B’). Beginning at stage III, however, keratin 10 became strongly localized to keratinocytes attached to the basement membrane and this pattern persisted through stage V (Fig. 4C’-D’). After metamorphosis, keratin 10 localized exclusively to the basal keratinocytes (Fig. 4F’). Taken together, our data shows that cell proliferation occurs evenly throughout the epidermis and is only restricted to basal keratinocytes after metamorphosis when the epidermis transforms into a stratified epithelium. Cryptic stratification of the epidermis actually began as early as stage III when the dermis began to stratify and keratin 10 localized to keratinocytes attached to the basement membrane.

**Figure 3.**
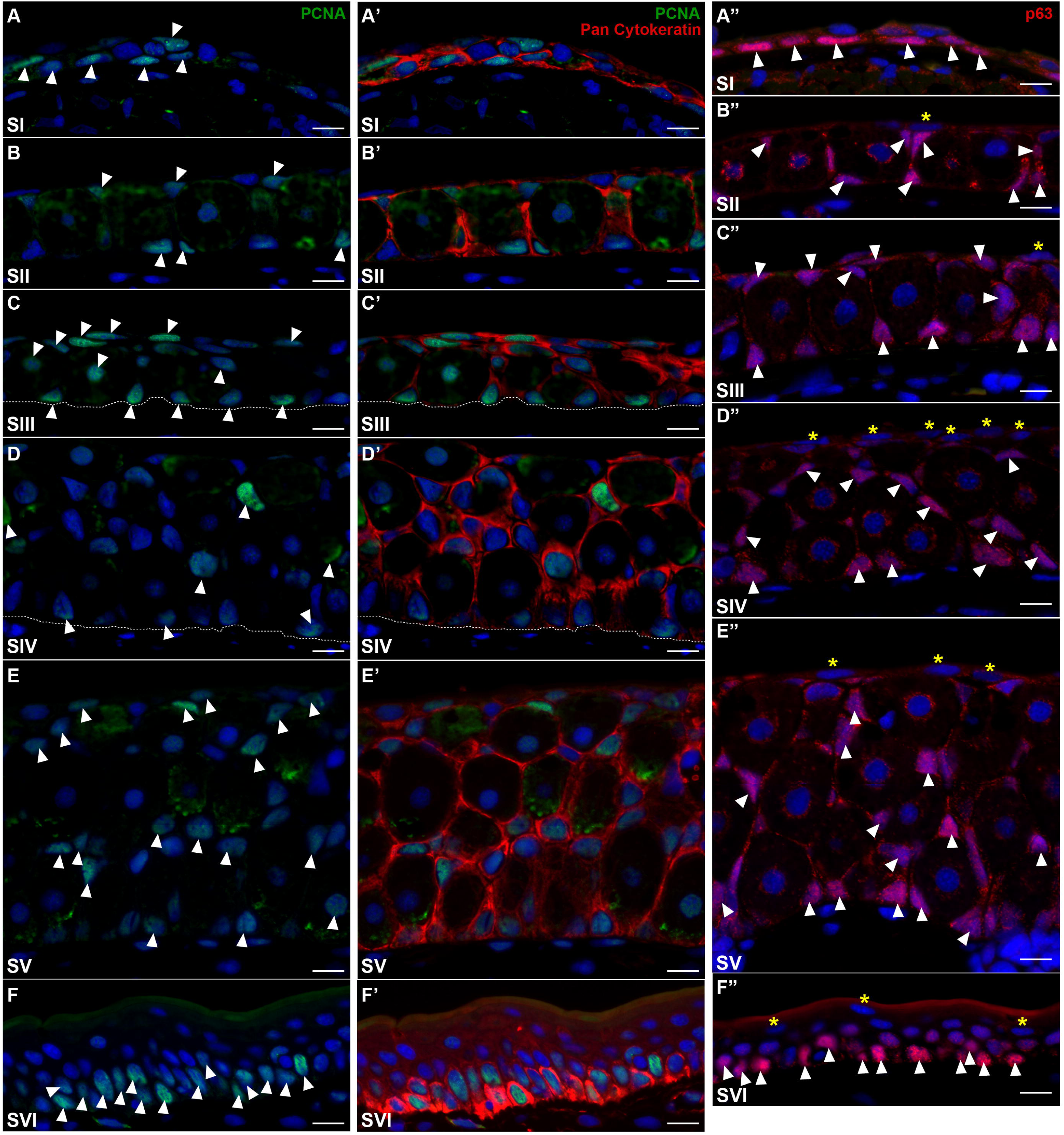
Keratinocyte proliferation is distributed throughout the epidermis and restricted to basal keratinocytes after metamorphosis. (A-E’) PCNA localization in the nucleus (white arrows) (A-F) demonstrates even distribution of cell proliferation within keratinocytes (A’-E’) throughout the epidermis of skin stages I-V. A pan-cytokeratin antibody was used to identify keratinocytes which also localized to Leydig cell membranes (A’-F’). Dotted white lines define the base of the epidermis (C-D’). (A”-E”) p63+ keratinocytes (white arrows) distributed throughout the epidermis similar to PCNA in skin stages I-V. p63 was absent in flattened keratinocytes of the stratum corneum (yellow asterisks). (F-F”) Following metamorphosis, PCNA and p63 were restricted to basal keratinocytes. Several suprabasal keratinocytes were weakly positive for both markers. Cytokeratin reactivity was also higher in basal compared ti suprabasal keratinocytes. PCNA = green, pan cytokeratin = red, Hoechst = blue (A-F’). p63 = red, Hoechst = blue (A”-F”). Scale bars = 20μm.

**Figure 4.**
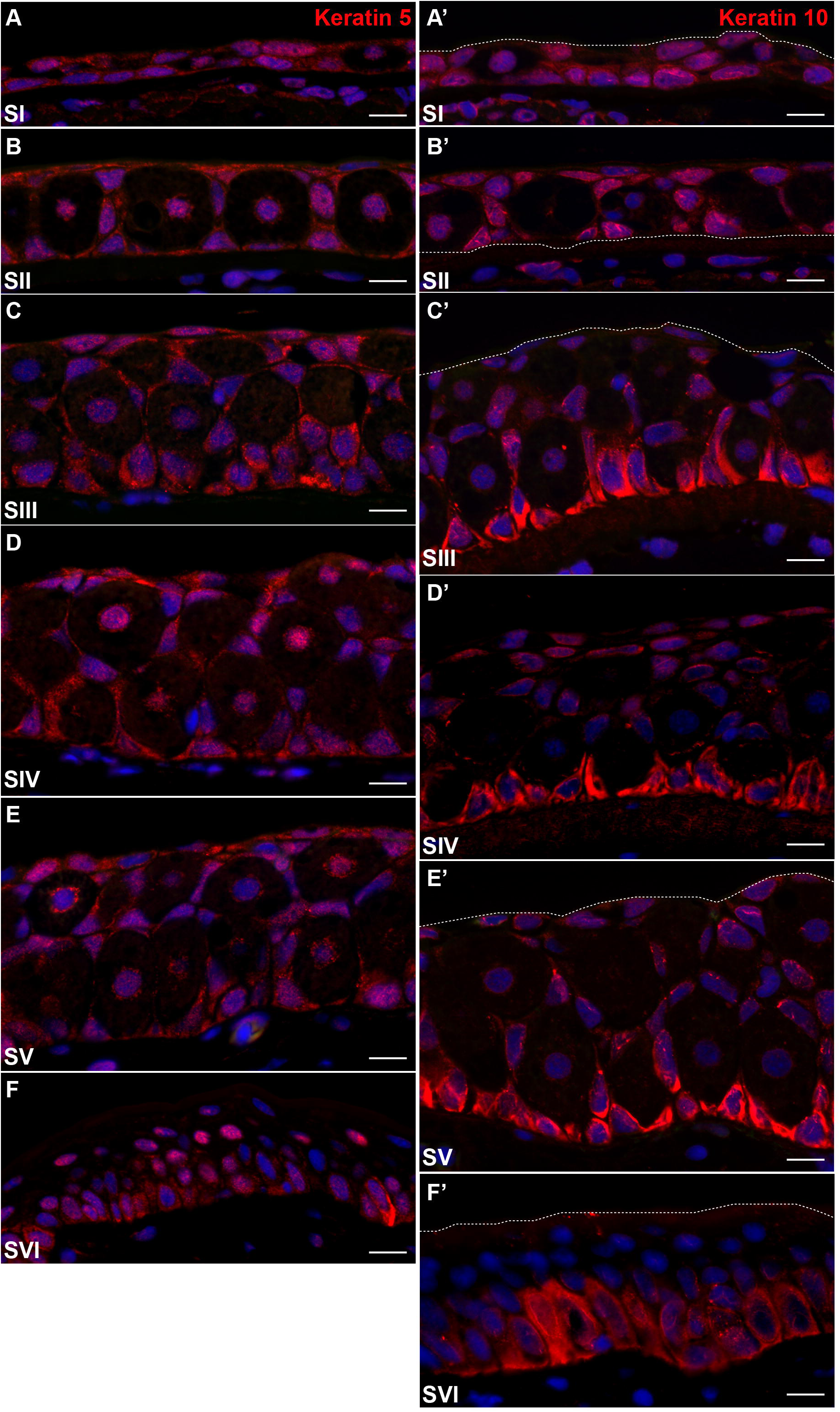
Keratin 10 localization reveals cryptic stratification of keratinocytes in developing axolotl epidermis. (A-E) Antibody to human keratin 5 localizes to all keratinocytes in developing epidermis through stage V. (F) Following metamorphosis keratin 5 localization is nuclear and cytoplasmic in basal keratinocytes and nuclear only in suprabasal keratinocytes. (A’-B’) Keratin 10 is ubiquitously expressed during stage I-II in all keratinocytes. (C’-E’) Beginning at stage III, keratin 10 is more strongly associated with basal keratinocytes, but remains associated with keratinocytes in all epidermal layers. (F’) After metamorphosis keratin 10 becomes exclusively associated with basal keratinocytes. White dotted lines show upper (A’, C’, E’, F’) and lower bounds of the epidermis (B’). Hoechst = blue (A-F’). Scale = 20um.

### The dermal lamella is distinct from the basement membrane and is the stratum compactum

Early embryonic development of the dermis in urodeles has remained obscure, especially in those species that remain aquatic and undergo cryptic metamorphosis (e.g., *Ambystoma mexicanum*, *Necturus maculosus*, etc.). Beginning in stage 2 skin we observed a thickened, acellular network of collagen beneath the epidermis (Fig. 1C-C’) similar to descriptions in larval salamander, frogs and fish where authors have variously referred to this layer as the basement lamella or collagen lamella (Hay and Revel, 1963; Kemp, 1959; Le Guellec et al., 2004; Takagi, 1956; Utoh et al., 2000; Weiss and Ferris, 1954). To better understand this collagen layer in axolotl skin, we used transmission electron microscopy (TEM) to further analyze its anatomy and association with the basement membrane (Fig. 5A-D). The thick collagen layer beneath the epidermis was immediately recognizable as the basement lamella with its network of orthogonally arranged collagen fibers bundled into plies (Fig. 5A-D). Upon close inspection, we observed hemidesmosomes linking the epidermis to the outermost layer of the basement lamella which was clearly distinct from the underlying collagen bundles and was the basement membrane (Fig. 5B-C). Thus, the basement membrane was directly occluded to the collagen fibers (Fig. 5C). In stage IV skin, fibroblasts and pigment cells were observed invading between these collagen plies and coincident with this invasion, we observed this collagen layer separating from the epidermis and basement membrane (Fig. 1D, E’ and Supplementary Fig. 1C). Because this collagen layer ultimately separates from the basement membrane to become the stratum compactum (lower layer of the dermis), we feel it is more appropriate to refer to this structure as the dermal lamella (Fig. 5A-D).

**Figure 5.**
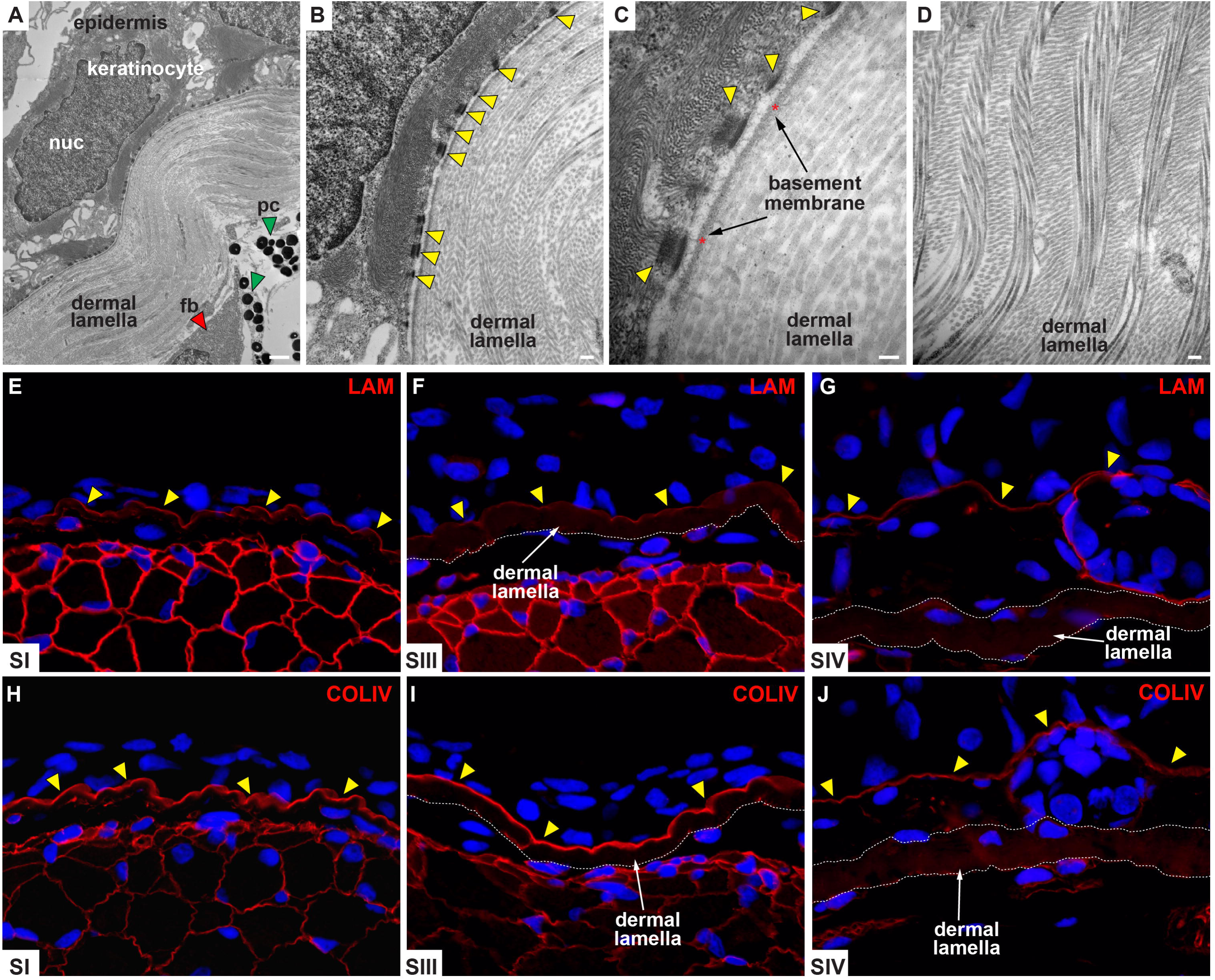
The dermal lamella is distinct from the basement membrane and is composed of orthogonally arranged collagen bundles. (A-D) Transmission electron microscopy (TEM) reveals direct association of basement membrane and dermal lamella. (A) Keratinocytes are visible above the basement membrane with fibroblasts (fb) and pigment cells (pc) beneath the dermal lamella. (B-C) Keratin fibers are visible attached to hemidesmosomes (yellow arrows) linking the epidermis to the basement membrane (red asterisks). (D) Collagen fibers comprising the dermal lamella are arranged into bundles that are aligned orthogonally to one another. (E-G) Laminin localization during axolotl skin development indicates lamina lucida component of the basement membrane (yellow arrows). The dermal lamella appears in stage III skin and is negative for laminin. (H-J) Collagen IV localization indicates lamina densa component of the basement membrane (yellow arrows). Similar to laminin localization, the dermal lamella is negative for collagen IV. Glands are encapsulated by a laminin+ membrane (G). Dermal lamella boundary is outlined by a white dashed line (E-J). nuc = nucleus.

In order to determine if the basement membrane was functionally distinct from the dermal lamella, we used localization of laminin and collagen IV to label the lamina lucida and lamina densa (Fig. 5E-F, H-I). Prior to formation of the dermal lamella (stage I-II), localization of laminin and collagen IV revealed that the mature basement membrane rested directly above the hypodermis (Fig. 5E, H). However, with the emergence of the dermal lamella at stage III, reactivity for these proteins revealed the basement membrane as a distinct band atop the dermal lamella (Fig. 5F, I and Suppl. Fig. 2). We also tracked separation of the basement membrane from the dermal lamella during skin development to determine if either of these matrix proteins contributed to the dermal lamella (Fig. 5E-J). As the dermal lamella separated from the basement membrane during dermal stratification (stage IV), the basement membrane remained affixed to the epidermis and the dermal lamella was negative for laminin and collagen IV (Fig. 5G, J). Together, our data demonstrates that as the dermal lamella forms and thickens, it remains distinct from the basement membrane and it is the separation of the two that facilitates subdivision of the dermis. Thus, invasion of fibroblasts and pigment cells into and on top of the dermal lamella, separation of the basement membrane from the top collagen ply of the dermal lamella and production of new connective tissue by invading fibroblasts during the transition from stage III-IV during skin leads to formation of the stratified dermis (stratum spongiosum and stratum compactum) (Fig. 1D-E and Fig. 5G, J).

### Delayed fibroblast infiltration reveals alternate trajectory for dermis regeneration that recapitulates development

With a clearer understanding of skin development in the axolotl, we turned our attention to investigating the degree to which skin regeneration might recapitulate the cellular events of skin development. In particular we were interested in how the stratum compactum reformed its complex orthogonal collagen plies. Previous work in salamanders and frogs showed that dermis regeneration proceeded through remodeling of newly deposited extracellular matrix (ECM) in the wound bed (Bertolotti et al., 2013; Levesque et al., 2010; Seifert and Maden, 2014; Seifert et al., 2012b; Yokoyama et al., 2011a). Analyzing full-thickness skin regeneration in light of our developmental data, it became apparent that dermis regeneration occurred uniquely from how the dermis developed (Fig. 6A-D and Suppl. Fig. 3). During healing, fibrosis occurred in the wound bed beginning approximately fourteen days post injury and by D21 the wound bed was filled with a dense ECM (Fig. 6A-B, I-I’ and Suppl. Fig. 3). After four weeks post-injury the wound bed ECM began remodeling into the denser stratum compactum and upper stratum spongiosum (Fig. 6C, J-J’). We observed the reorganization of the stratum compactum from wound bed matrix near the undamaged collagen fibers at the wound margins with increased remodeling occurring over time from the out margins to the center of the wound (Suppl. Fig. 3). At no time did we observe a dermal lamella forming in association with the epidermis during normal skin regeneration (Fig. 6C-D, I-K’ and Suppl. Fig. 3).

**Figure 6.**
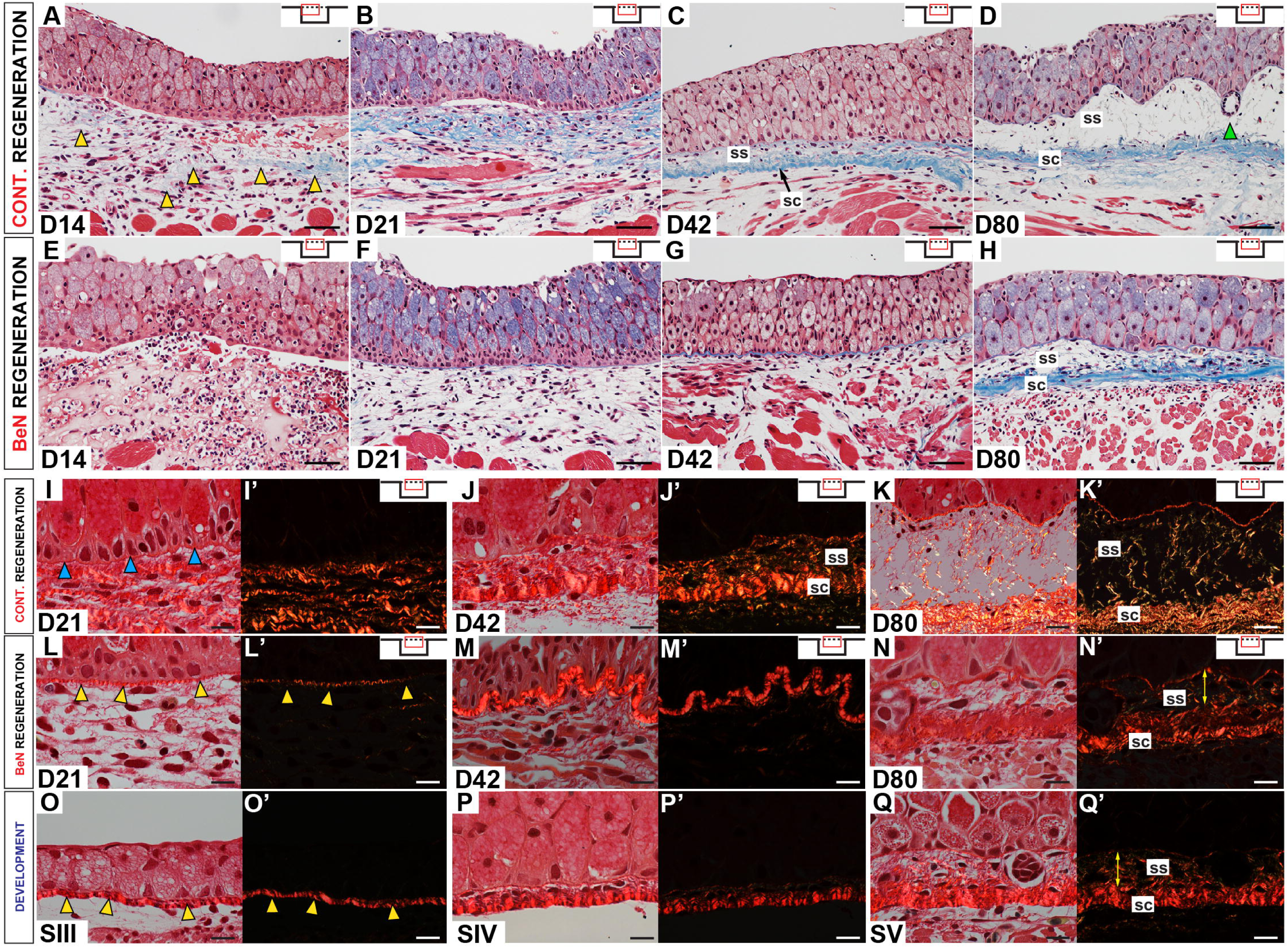
Inhibiting fibroblast migration into full-thickness skin wounds using beryllium nitrate reveals developmental pathway for dermis regeneration. (A-D) Full-thickness healing skin stained with Masson’s trichrome as it progresses through normal regeneration (n=5/timepoint). Extracellular matrix (ECM) (blue staining, yellow arrows) appears in the wound bed 14 days post injury (D14) (A). New ECM occupies the entire wound bed at D21 (B) and begins remodeling into a new stratum compactum by D42 (C). The result is regenerated skin, including a bi-layered dermis with new glands (green arrow) by D80 (D). (E-H) Full-thickness skin wounds treated with beryllium nitrate (BeN) undergo delayed healing and regeneration (n=5/timepoint). There is almost no ECM in the wound bed except in association with the epidermis beginning at D21 (F-G). At D80 the dermis is regenerating and a stratum spongiosum (ss) and stratum compactum (sc) are visible. Full-thickness skin stained with picrosirius red during normal regeneration shows a collagen-negative basement membrane at D21 (blue arrows) (I) and collagen deposition in the wound bed (I’) that begins to subdivide by D42 (J-J’). Fully regenerated skin at D80 showing a collagen-rich basement membrane, thin collagen fibers of the stratum spongiosum, and dense collagen fibers in the stratum compactum (K-K’). (E-H) BeN-treated skin wounds show little to no collagen in the wound bed except for the dermal lamella in association with the epidermis (L-M’). A thin band of collagen appears at D21 (yellow-arrows) (F, L-L’) and thickens into a much denser sheet by D42 (M-M’). After eighty days the dermal lamella separates from the basement membrane as the stratum spongiosum (ss) and stratum compactum (sc) begin to form (yellow arrows) (N-N’). Healing BeN-treated wounds mirror the progression of skin development as immature stage I skin also forms a dermal lamella that begins as a thin band collagen matrix that thickens through stage III (O-O’). Fibroblasts invade the dermal lamella during development (P, P’) and the dermis becomes stratified in mature skin (yellow arrows) (Q-Q’). Insets in A-N’ show position in wound bed. SS = stratum spongiosum, SC = stratum compactum. Scale = 50μm (A-H), 20μm (I-Q, I’-Q’)

Previous work from our lab showed that treating dorsal skin wounds with beryllium nitrate (BeN) inhibited local fibroblast migration into the wound bed and this delayed skin regeneration (Cook and Seifert, 2016). Analyzing BeN-treated skin wounds (n=5/timepoint), we observed very little matrix production in skin wounds even after six weeks post injury (Fig. 6E-G and Suppl. Fig. 3). The result was that the epidermis remained relatively close to the underlying skeletal muscle. Although we observed a paucity of new matrix in the wound bed, we did observe a collagen-rich layer in association with the epidermis that was structurally similar to the dermal lamella (Fig. 6G, L-M’ and Suppl. Fig. 3). In fact, healing BeN-treated wounds recapitulated the cellular and extracellular events we observed during skin development almost perfectly (Fig. 6L-Q’ and Suppl. Fig. 3). During both events, the dermal lamella emerged in association with the epidermis and thickened over time (Fig. 6L-L’, O-O’, M-M’). In BeN-treated wounds, fibroblasts from outside the wound margins eventually make their way into the wound bed (Cook and Seifert, 2016) and after eighty-days we observed fibroblasts invading into and above the dermal lamella (Fig. 6N-N’). These fibroblasts produced new connective tissue that regenerated the stratum spongiosum and continued separation of the dermal lamella from the basement membrane produced the stratum compactum (Fig. 6N-N’). This sequence of events mirrored the transition from stage III-V during skin development (Fig. 6P-Q’). These data demonstrate that normal dermis regeneration in salamanders occurs through remodeling of wound bed ECM, instead of through production and subdivision of the dermal lamella. However, our data also shows that preventing injury-induced fibroblast migration and matrix deposition in the wound bed can elicit a developmental pattern of dermis formation. This result demonstrates that salamander dermis can regenerate through two different mechanisms: one that relies on matrix remodeling and one that produces dermis *de novo* as occurs during development.

## Discussion

In an effort to characterize epidermal and dermal development in salamanders to consider the extent to which regeneration recapitulates developmental processes, we analyzed post-embryonic development of full-thickness axolotl skin (epidermis and dermis) and found that it proceeded through six recognizable stages (I-VI) where stage V represents the juvenile/adult morphology in normally paedomorphic animals and stage VI represents the penultimate stage that occurs after induced metamorphosis. Throughout stage I of skin development Leydig cells began to populate the epidermis whereas during stage II, Leydig cells occupied most of the epidermis, melanocytes appeared beneath the basement membrane, and the dermal lamella began to form in association with the epidermis. The transition to stage III was marked by a thickening of the epidermis and dermal lamella, and keratinocytes aggregated to form nascent glands. The most prominent transformation of the skin occurred between stage III-IV when the bi-layered dermis began to form, and keratin 10 expression revealed a cryptic stratification of keratinocytes in the epidermis. Independent of this stratification, keratinocyte proliferation occurred throughout the epidermis and was not restricted to basal keratinocytes until after metamorphosis when the epidermis transitioned to a fully stratified epithelium. During formation of the dermis, the dermal lamella delaminated from the basement membrane as dermal fibroblasts infiltrated through the stacks of collagen fibers and began producing new connective tissue of the stratum spongiosum. In adult paedomorphic skin, the stratum spongiosum contained loosely organized collagen fibers, mucus and granular glands, melanocytes and fibroblasts. The dermal lamella, which becomes the stratum compactum, retained its morphology of orthogonally arranged collagen fibers and remained largely acellular. Following metamorphosis, Leydig cells completely disappeared from the epidermis as it differentiated into a fully stratified epithelium. In response to thyroxine, granular glands increased in size to occupy most of the stratum spongiosum while the overall arrangement of the dermis did not change (Seifert et al., 2012b).

In determining these stages of skin development, we discovered that progression through each stage is a function of animal size. Under highly controlled conditions where post-hatching animals were raised in isolation, mass, length and age were all strong predictors for a particular stage of skin development. However, when animals were raised in groups, a situation more akin to normal laboratory and wild populations, age was only 18% effective in accurately predicting the stage of skin development whereas mass or total length could predict skin stages in 90% of selected animals. While density-dependent effects on growth rate in salamanders are well known (Semlitsch, 1987), our work demonstrates that density can directly impact developmental rate for a particular tissue, a notion theorized by other investigators modeling a relationship between body size and developmental staging (Gillooly et al., 2002; Gillooly et al., 2007). The variability in developmental progression we observed in axolotls is similar to variation reported for frog skin development where limb size, but not post-hatching age, was the best predictor of skin stage prior to metamorphic climax (Kawai et al., 1994). This pre-metamorphic variability in developmental progression disappeared as rising levels of serum thyroid hormone reached peak levels (Kawai et al., 1994; Tamakoshi et al., 1998) supporting the idea that thyroid hormone levels are a major driver of skin development in frogs. While this may be the case for anurans, axolotls are neotenic and have uncoupled thyroid hormone driven metamorphosis from normal post-embryonic development (Brown, 1997; Rosenkilde and Ussing, 2004). Thus, the mechanisms tying body size to skin development in salamanders awaits further investigation as does whether rising thyroid hormone levels are tied to body size or growth rate. Future studies targeting genetic or hormonal indicators of cryptic metamorphosis may shed light on how skin development is regulated in neotenic salamanders.

Because axolotls are neotenic, they retain certain larval characteristics in the sexually mature adult form. One such larval feature is the presence of Leydig cells in the epidermis. The persistence of these cells throughout life is an enigmatic and unique feature of paedomorphic salamanders as these cells normally disappear during metamorphosis in terrestrial urodeles (Jarial, 1989; Kelly, 1966; Ohmura and Wakahara, 1998; Seifert et al., 2012b). A similar phenomenon occurs in anurans where skein cells are the dominant epidermal cells type in larval skin, but are lost during metamorphosis in response to increasing levels of thyroid hormone (Izutsu et al., 1993; Kawai et al., 1994; Robinson and Heintzelman, 1987). While Leydig cells undergo a distinctive maturation that is defined by an increasing density of granules produced from the nucleus (Gerling et al., 2012; Kelly, 1966), whether this is autonomous or dependent on molecular cues from keratinocytes remains unknown. Some authors have speculated that Leydig cells contain anti-microbial peptides (Jarial, 1989) while others have suggested they mediate key signals necessary for regeneration (Kumar et al., 2010). Whatever their true function, they are the most prominent feature of axolotl epidermis. Further studies should focus on antimicrobial and antifungal properties of these cells including the biochemical nature and contents of the granules.

Development of the vertebrate epidermis is generally defined by a progressive differentiation program whereby a simple layer of ectodermal cells transforms into a fully stratified epithelium comprised of a basal germinative layer, suprabasal granular/spinous layer and an outer periderm layer that is ultimately replaced with a cornified barrier layer (Koster and Roop, 2007). Structural analysis of frog and fish skin suggests that differentiation of the epidermis may have evolved for a terrestrial lifestyle as fish epidermis lacks stratification (Henrikson and Matoltsy, 1967) and frog skin acquires a hierarchical stratification only after metamorphosis (Izutsu et al., 1993; Kawai et al., 1994; Robinson and Heintzelman, 1987). Our cellular and molecular data demonstrates that axolotl epidermis up to stage III is similar to anuran epidermis prior to metamorphosis (Kinoshita and Sasaki, 1994; Tamakoshi et al., 1998). Similar to frogs, we found proliferating (PCNA+) cells distributed evenly throughout the entire epidermis and all keratinocytes were positive for keratin 5 and 10 supporting the apparent lack of epidermal stratification during early skin development. As larval anurans approach metamorphic climax, fibroblasts invade the dermal lamella and secrete proteases that help separate it from the basement membrane (Berry et al., 1998; Izutsu et al., 1993; Tamakoshi et al., 1998). Simultaneously, multiple epidermal lineages disappear leaving basal keratinocytes that express Rana adult keratin (RAK) to populate the adult epidermis (Suzuki et al., 2002). In axolotl skin, the transition from stage III-IV resembles maturation of frog skin at metamorphic climax where fibroblasts invade the dermal lamella as it separates from the basement membrane (Izutsu et al., 1993; Tamakoshi et al., 1998). Coincident with this transition we found that an antibody to human keratin 10 strongly labeled keratinocytes attached to the basement membrane and to a lesser extent suprabasal keratinocytes. This localization persists in adult axolotl skin and in combination with our histological data, shows that keratinocytes in contact with the basement membrane are distinct from suprabasal keratinocytes that express low levels of keratin 10.

We also investigated localization of p63 which is required for epidermal specification and stratification during development (Byrne et al., 1994; Lee and Kimelman, 2002). After being activated in surface ectoderm, p63 activates keratins 5 and 14 via AP-2 (Byrne et al., 1994; Koster et al., 2006) which helps to establish the germinative layer of basal keratinocytes. Although p63 appeared to localize to basal and suprabasal keratinocytes, in contrast to PCNA we found that it was largely absent from flattened keratinocytes in the stratum corneum. Thus, our data suggests that at least three types of keratinocytes (in addition to Leydig cells) populate adult axolotl skin: basal keratinocytes attached to the basement membrane that are strongly K10+, suprabasal keratinocytes that are weakly K10+ and flattened keratinocytes covering the epidermis that are p63-. Following induced metamorphosis and the loss of Leydig cells, a completely stratified epithelium emerges with a basal germinative layer, granular/spinous layer and stratum corneum. Proliferation was restricted to p63+ basal keratinocytes, and differentiation occurred towards the surface. It is important to note that axolotl keratins 5 and 10 show X% and X% similarity to human keratins 5 and 15 respectively suggesting that the keratins recognized by antibodies used in this study could be different keratin proteins. As genomic resources for the axolotl progress, better resolution of keratin protein structure will allow a more detailed examination of their expression pattern and function during development.

In conjunction with epidermal stratification we observed formation of a dermal lamella and its delamination from the basement membrane to form the stratum compactum. During formation of the dermal lamella, the organization of collagen fibers mirrored observations in anurans and fish (Fox and Whitear, 1990; Hay and Revel, 1963; Kemp, 1961; Kemp, 1963; Le Guellec et al., 2004; Salpeter and Singer, 1959; Tamakoshi et al., 1998; Utoh et al., 2000; Weiss and Ferris, 1954). Our TEM data supports previous work showing that the epidermis is the source of the dermal lamella (Hay and Revel, 1963) as does our data from BeN-treated skin wounds showing formation of a dermal lamella in close association with the epidermis when fibroblasts are absent from the wound bed (Cook and Seifert, 2016). Maturation of the dermis occurred when fibroblasts invaded the dermal lamella and it separated from the epidermis. The result was the production of new connective tissue and subdivision of the dermis into the upper stratum spongiosum and lower stratum compactum. Previous work in anurans showed that a similar fibroblast invasion of the dermal lamella was coincident with maturation of the dermis (Kemp, 1961; Kemp, 1963; Utoh et al., 2000) and that this event was regulated by thyroid hormone at metamorphic climax (Kawai et al., 1994). The activation of downstream target genes by thyroid hormone is specific to the epidermis and sub-epidermal fibroblasts where up-regulation of caspase-3 induces apoptosis of larval keratinocytes in the epidermis and induction of collagenase-3 in fibroblasts helps to separate the dermal lamella from the basement membrane (Schreiber and Brown, 2003). That fibroblasts appear to perform a similar function during axolotl skin development suggests that either activation of proteolytic enzymes is independent of thyroid hormone or that fibroblasts and keratinocytes are differentially sensitive to very low circulating levels of thyroid hormone in axolotls (Brown, 1997; Rosenkilde and Ussing, 2004). Further studies of this phenomenon that specifically investigate thyroid hormone sensitivity and cell-specific expression of thyroid hormone receptors will shed light on how metamorphosis and skin development can uncouple in neotenic animals.

Our investigation of skin development allowed us to examine the degree to which skin regeneration recapitulated developmental events or displayed a unique regenerative program. Comparing skin regeneration to development, we discovered that the rebuilt dermis is remodeled directly from wound bed ECM and not by generating an epidermally-produced dermal lamella. By using BeN to prevent fibroblast migration into the wound bed (Cook and Seifert, 2016) we also discovered that the epidermis *can* generate a dermal lamella and thus dermis regeneration can occur by a secondary mechanism that recapitulates development. Previous studies of axolotl skin regeneration have shown that re-epithelialization occurs 24hrs after injury with keratinocytes migrating to cover the wound bed (Levesque et al., 2010; Seifert et al., 2012b). During the first two weeks post injury in adult animals the collagen IV+ lamina densa and orthogonally arranged collagen fibers of the dermal lamella are absent from the wound bed (Cook and Seifert, 2016; Seifert et al., 2012b). Approximately two weeks post injury new extracellular matrix is deposited in the wound with tenascin-C, fibronectin, and hyaluronate dominating the wound bed. As regeneration proceeds, collagen is slowly deposited, the mature basement membrane is replaced and the stratified dermis forms *de novo* from remodeled wound collagen (Seifert et al., 2012b). Remodeling of the stratum compactum occurs from the wound margins to the center and continues over the next one hundred days. In contrast, the stratum compactum first appears as the dermal lamella during development and is produced by the epidermis (Hay and Revel, 1963). It is only after fibroblasts invade the dermal lamella that the stratum spongiosum begins to form. When we prevented fibroblasts from entering the wound site using beryllium nitrate, the wound remained free of extracellular matrix until fibroblasts returned (Cook and Seifert, 2016). Prior to this reinvasion of the wound bed with fibroblasts we observed formation of the dermal lamella beneath the epidermis. Approximately two months post injury we observed separation of the dermal lamella from the basement membrane as glands descended from the epidermis. This series of events paralleled our observations of skin development and demonstrates that axolotl skin can regenerate along two different routes: one that involves fibrosis and remodeling of the ECM and another that recapitulates the events of skin development. Previous work wounding larval salamander skin found that when wounds were created prior to separation of the dermal lamella, regeneration proceeded akin to development (Hay and Revel, 1963; Weiss and Ferris, 1956). These data support the idea that fibroblast-driven fibrosis in skin wounds is an ancient repair mechanism (Seifert and Maden, 2014) and that regenerating organisms have the ability to appropriately remodel the local ECM whereas non-regenerating animals are unable to appropriately reconstruct the original dermal architecture and are left with scar tissue.

## METHODS

### Animals

Axolotls (*Ambystoma mexicanum*) were acquired from our own laboratory colony or from the Ambystoma Genetic Stock Center (Lexington, KY) and maintained between 17-18°C in modified Holtfreter’s Solution. Larval animals were fed brine shrimp until they could eat California blackworms (*Lumbriculus variegatus*, J.F. Enterprises) and received approximately the same amount of food at each feeding. To generate growth curves and to establish stages of skin development, sibling animals from the same clutch were reared individually. Individual housing excluded any effects of density on growth and development. Additional batches of sibling animals were group-housed to address differences in growth due to density. Prior to tissue harvest axolotls were anesthetized by full submersion in 0.01% (aqueous) Benzocaine (Sigma). Metamorphic animals were also used for this study. Metamorphosis was induced with L-thyroxine as detailed previously (Coots & Seifert, 2015). All procedures were conducted in accordance with, and approved by, the University of Kentucky Institutional Animal Care and Use Committee (IACUC Protocol: 2013-1174).

### Full-thickness excisional (FTE) wounding and beryllium nitrate (BeN) treatment

Axolotls were anesthetized by full submersion in 0.01% (aqueous) Benzocaine (Sigma). 4mm biopsy punches (Sklar Instruments, West Chester, PA) were used to create full-thickness excisional (FTE) wounds through the skin into the dorsal muscle. Two dorsal skin wounds were created posterior to the forelimbs and anterior to the hindlimbs. Application of beryllium nitrate treatment was carried out as previously described (Cook and Seifert, 2016). Briefly, 100 mM beryllium nitrate solutions were prepared by diluting a stock solution (35% w/v) (Sigma) in distilled water. Following an FTE wound, animals were submersed in 100 mM BeN for 2 minutes, after which they were continually rinsed in running tap water for 10 minutes. After rinsing, a second wound corresponding to the mirror position across the dorsal midline was made. Each animal served as its own control. Healing wound tissue was harvested at indicated days post injury and prepared for paraffin embedding.

### Histology

Freshly harvested tissues were fixed overnight (~16-24hr) in 10% neutral buffered formalin (NBF) (American Master Tech Scientific Inc.), washed in phosphate buffered saline (PBS) 3 times, dehydrated in 70% EtOH and processed for paraffin embedding using a microwave tissue processor (Histos5). Tissue samples were sectioned at 5μm. Tissue sections were stained with Masson’s trichrome (Richard-Allen Scientific) or Picrosirius Red (American Master Tech) as previously described (Seifert et al., 2012).

### Growth Curve and Skin Development Staging

We raised two sibling cohorts to generate growth curves and to develop a normal staging series to study skin development. Animals from cohort one (n=20) were used to ensure a standard growth curve at 17-18°C using our feeding regimen and animals from cohort two (n=31), which was treated identically to cohort one, were used to characterize skin development using histological analysis. Cohort two consisted of two batches of animals so as to completely cover the size range that captured all skin stages. Cohorts were weighed (g) and measured (total body length - cm) every seven days. Animals (n=3) were sampled at random every seven days. For smaller animals the entire animal was prepared for sectioning and for larger animals we prepared full-thickness dorsal skin. Key cellular characteristics were used to determine skin development stages. We used the following criteria to determine whether a cellular or morphological event was diagnostic for a particular stage: a) the cellular event must be visually distinct and recognizable using standard histological analysis, b) the event must be consistently present within the sampled population, and c) the event must be consistent in its chronological appearance. In those cases where we were unable to separate cellular events in time, these events would be grouped together to characterize a single stage.

### Immunohistochemistry

For immunohistochemistry, slides were de-paraffinized, rehydrated and antigen retrieval was performed. Sections were washed in tris-buffered saline (TBS), blocked for biotin and avidin, (Vector Laboratories Inc., Burlingame, CA), and incubated with 1° antibodies overnight at 4°C. Isotyped IgGs were used as negative controls and run at matching concentrations to the primary antibody. Following incubation, slides were washed in TBS and incubated with biotinylated 2° antibody (Vector) for 30 minutes. Antibodies were visualized using streptavidin conjugated Alexa-Flour 488 or 594 (Invitrogen, Carlsbad, CA). Nuclei were counterstained with either 10μg/ml Hoescht (Invitrogen) for fluorescence or working Weigert’s hematoxylin (Thermo Fisher Scientific, Waltham, MA) for bright field visualization.

Antibodies used were as follows: Collagen IV (Rockland Inc., Limerick, PA) 1:450, Collagen I (Rockland Inc.) 1:500, Keratin 5 and 10 (BioLegend, San Diego, CA) 1:500, Pan-Cytokeratin (Agilent, Santa Clara, CA) 1:500, PCNA (Agilent) 1:1000, Laminin (Agilent) 1:750, p63 (Nordic MUbio, Susteren, Netherlands) 1X, and axolotl Fibronectin (gift of T. Darribere) 1:300. For antigen retrieval on paraffin sections we used the microwave component of our Histo5 with citrate buffer (pH 6.0) for 25 minutes at 85°C.

### Microscopy and image acquisition

Bright-field images were taken on a BX53 light microscope (Olympus) using a DP80 CCD camera (Olympus). Whole mount images were taken on an SZX10 light microscope (Olympus) using a DP73 CCD camera (Olympus). A dual mCherry-GFP filter was used to detect auto-fluorescing erythrocytes.

### Transmission Electron Microscopy

Harvested tissue was fixed in 4% paraformaldehyde with 2-3.5% glutaraldehyde in 0.1M Sorenson’s phosphate buffer (1.5hr at 4 degrees C), washed with 5% sucrose in Sorenson’s and fixed again in 1% OsO4 in 0.1M Sorenson’s, and dehydrated in 50-100% ethanol for 48hrs. Samples were infiltrated with propylene oxide for 30min prior to resin embedding. Resin embedding used 50/50 resin/propylene oxide with an accelerator and samples were left under a 60-watt lamp for 1hr. This was repeated for 100% resin. Tissue samples were thin sectioned using a Reichert Ultracut E UltraMicrotome.

### Statistics

Data analysis was performed using SPSS 22.0 for Windows (IBM SPSS Software, Armonk, New York, USA) and JMP 12 (SAS, Cary, NC). Linear regression was used to test organismal characters (age, length, and weight) as predictors of developmental stage. Because the relationship between mass and stage was an exponential function we log-transformed the data by a factor of 2 for the regression analysis. The weight and length predictor variables were measured experimentally and the response variable (stage of skin development) was determined through histological analysis of the stained tissue. Data used to generate the regressions is provided in Supplementary Data Table 1.

## Acknowledgements

We would like to thank Katherine Thompson for statistical consultation, members of the Seifert lab for insightful discussions and three anonymous reviewers for constructive comments. We also thank Laura and Chris Muzinic of the Ambystoma Genetic Stock Center at the University of Kentucky for supplying some of the embryos used in this study.

## Author Contributions

D.S, A.B.C and A.W.S developed the project, designed and carried out the experiments. A.B.C and A.W.S wrote and revised the manuscript.

