## Supplemental Figures for "Inhibiting fibroblast aggregation in skin wounds unlocks developmental pathway to regeneration"

Supplementary Figures

**Supplementary Figure 1. Histology of ampullae and glands within developing axolotl skin.** (A-C) Ampullae stained with Masson's trichrome (green arrows). (D) Cells forming ampullae showed reactivity to pan-cytokeratin antibody (green arrow). (E) Glands formed from keratinocytes (yellow arrow). Scale = 10um (A), 20um (B, C).

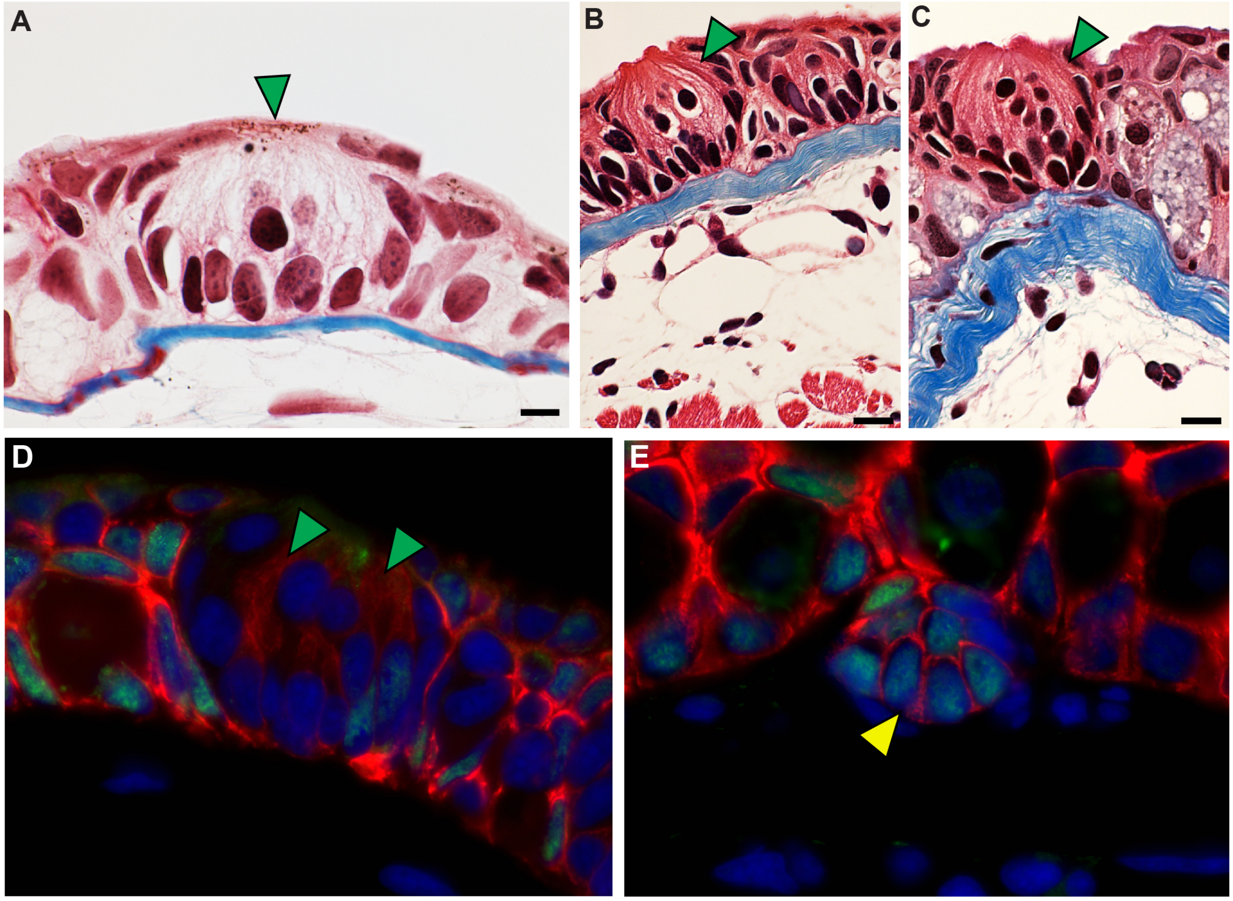

**Supplementary Figure 2. The dermal lamella is composed of collagen.**

Stage II and IV developing axolotl skin stained with an antibody for collagen I. Scale = 20um.

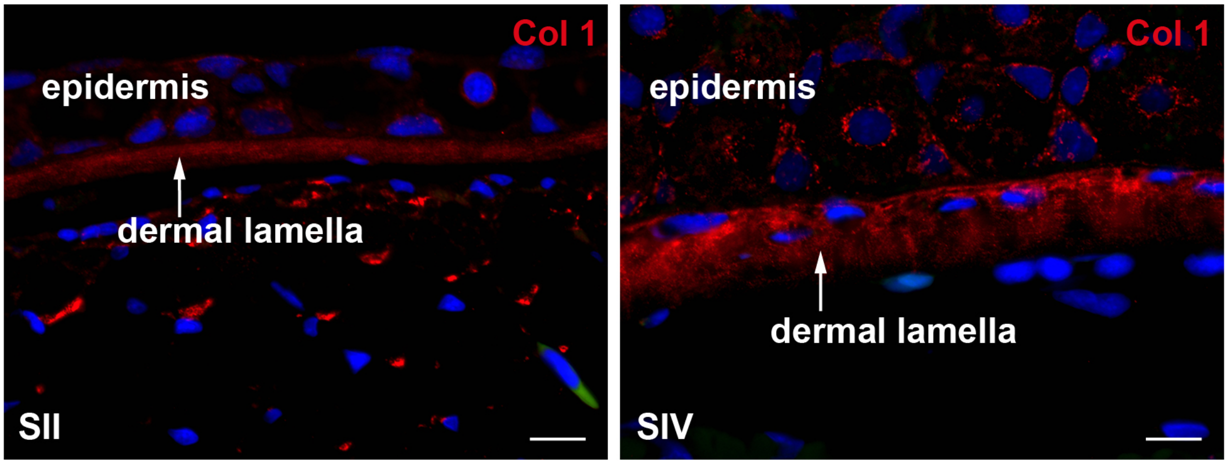

**Supplementary Figure 3. Wound bed detail of regenerating skin, regenerating skin treated with beryllium nitrate and developing skin.** During normal skin regeneration the wound bed is filled with new extracellular matrix. This matrix is remodeled into the stratum compactum (sc) and stratum spongiosum (ss). Regenerating skin treated with beryllium nitrate (BeN) forms a thin band of extracellular matrix connected to the basal layer of the epidermis by D21 (yellow arrows). The newly formed dermal lamella (dl) thickens by D42. The basal lamina separates from the dermal lamella prior to D80. Normal skin development follows this pattern closely (SI-SV). Scale bars = 50µm (Cont. Regen. D80, D139), 20µm (Cont. Regen. D21, 42; BeN Regen. D80; Development SI-V).

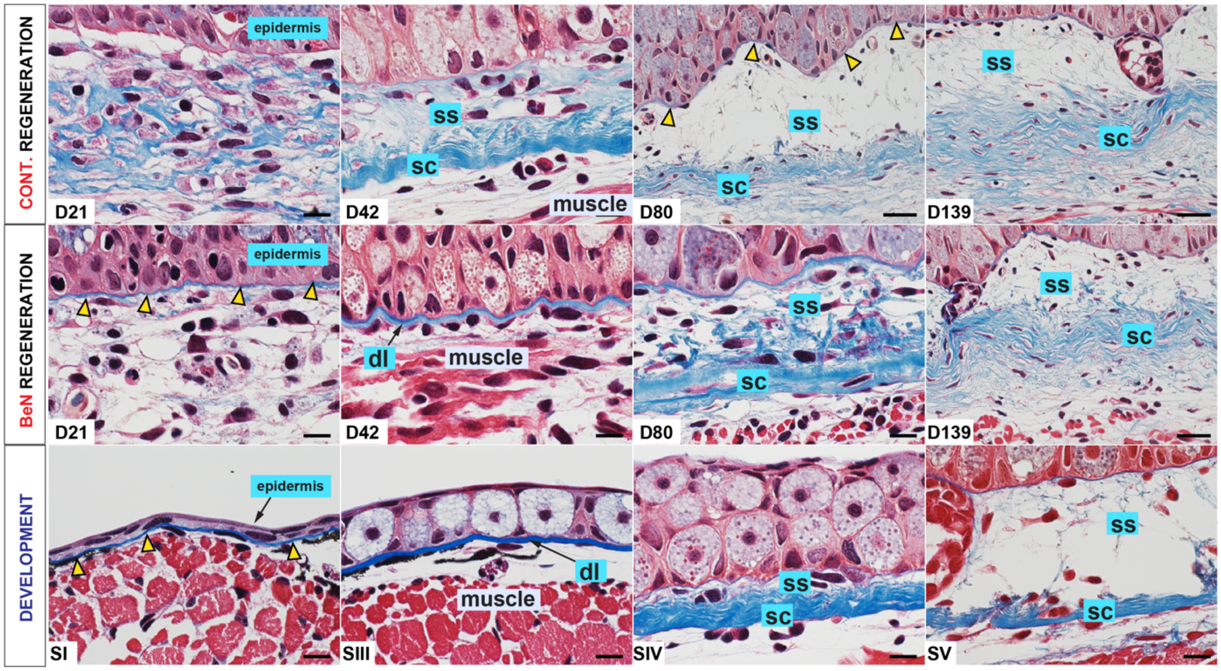
